# Mechanical characterization of compliant neural probes, their insertion regimes and a rule of thumb design

**DOI:** 10.1101/770107

**Authors:** Zac Smith

## Abstract

Investigates mechanical theory of buckling compliant neural probes, how this relates to insertion forces using normalized force data and discusses potential strategies to improve the insertion ability of neural probes. This report also provides a guide for mechanically testing compliant neural probes and demonstrates a design rule of thumb that insertion forces should be 1/3 lower than the buckling force to enable reliable insertion.

## Introduction

Compliant probes show promise in improving the long-term performance of *in vivo* neural probes, who’s current rigid materialled designs induce an immune response in tested subjects[1]–[6]. However, compliant neural probes have an array of barriers to overcome to be competitive with traditionally rigid neural probes. One major issue is the reliability of insertion into brain tissue. Compliant polymer probes for example have a much lower Youngs Modulus than silicon, is less stiff and less able to pierce the brain. This constrains the depth to which compliant neural probes can reach into the brain, limiting most current studies to near surface regions. Alternative methods are being employed to strengthen these probes, such as being encased in a rigid sheath which is withdrawn once inserted. However, these techniques likely add to the trauma of the surgical procedure, negating the gains in switching to compliant probes. This paper brings together some of the theoretical reasons why compliant probes behave as they do and combines it with mechanical testing which would serve as a platform to inform on better design and techniques for compliant neural probes.

## Theory

### Regimes of neural insertion

A single neural probe can be modelled as a thin, uniform beam, with one end fixed and the other end pinned against the brain’s surface using Euler’s buckling formula (Eq. 1)[7]. A ubiquitous problem for compliant materialled probes is their increased risk of mechanical buckling during the insertion process. On insertion of a single shank probe into tissue, the result can be one of three:

1. **full insertion** of the probe into the desired tissue, no buckling behavior (*F*_*Euler*_ ≫ *F*_*Insertion*_),
2. **buckling** on contact with the tissue surface with no apparent insertion (*F*_*Euler*_ ≤ *F*_*Insertion*_),
3. **partial insertion** before buckling (*F*_*Euler*_ ≫ *F*_*Insertion*_ | *F*_*Euler*_ ≤ *F*_*shear*_).

For each regime, the forces of import and their relationships are displayed. The probe-tissue system generally prefers the regime induced by the lower force. Three forces primarily govern whether buckling or insertion occurs: Euler’s buckling force (*F*_*Euler*_), the insertion force (*F*_*Insertion*_) and the shear force (*F*_*shear*_). Buckling most likely occurs because of regime 2 rather than three, because the force required to buckle the probe increases as the probe is inserted. Euler’s buckling force is a compressive force applied axially on a probe that causes a large deflection of the probes profile and is expressed by Euler’s formulae:

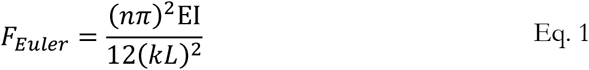

This formula is often used for compliant probes in the literature [7], [8] where *n* is the mode of buckling, generally taken as 1 unless this mode is artificially restricted, E is the Young’s Modulus of the probe’s material, I the second moment of area, L the probes length and k the probes effective length. The value for k varies from 0.5 – 2 depending on the boundary condition of the probe-tissue contact [9]–[11]. The column effective length factor *k* captures the degree to which each end of the column is constrained against movement. For a probe inserting into brain tissue, the base of the probe is considered to be clamped to the insertion tool and fixed (allowing for no translation or rotation), and the tip of the probe is pinned in the x-y plane as soon as it contacts brain tissue (only allows for rotation, not translation). Thus, the commonly accepted value of k is 0.7, validated experimentally in [12].

### Slipping phenomenon

The major issue during testing is the potential slipping of shanks before buckling on the metal plate. This was solved in this investigation by lowering the speed of the headstand for buckling tests, since faster speeds more likely resulted in slipping before buckling could be measured. Shank slipping can be viewed as an alternative to the three regimes and is characterized by a loss of loading force. An example of slipping occurring after buckling can be seen below for two-probe buckling.

As shown in the Figure 1, during pre-contact the probes are advanced by an actuator at a constant speed towards a perpendicular metal plate. On contact with the metal surface, the probes withstand some loading before buckling. Buckling represents a flatlining of the force vs distance curve, since the shanks do not withstand any continued loading, the amplitude of the displacement increases instead as the headstand compresses the probes. A force drop after buckling is characteristic of a shank slipping on the metal plate. Since the probes amplitude of displacement is increasing with distance, the tip of the probe is making a shallower angle to the plane of the metal plate, increasing the chance of slip. After all shanks have slipped, a non-zero force remains representing the two slipped probes bending at its new slipped location.

**Figure 1:**
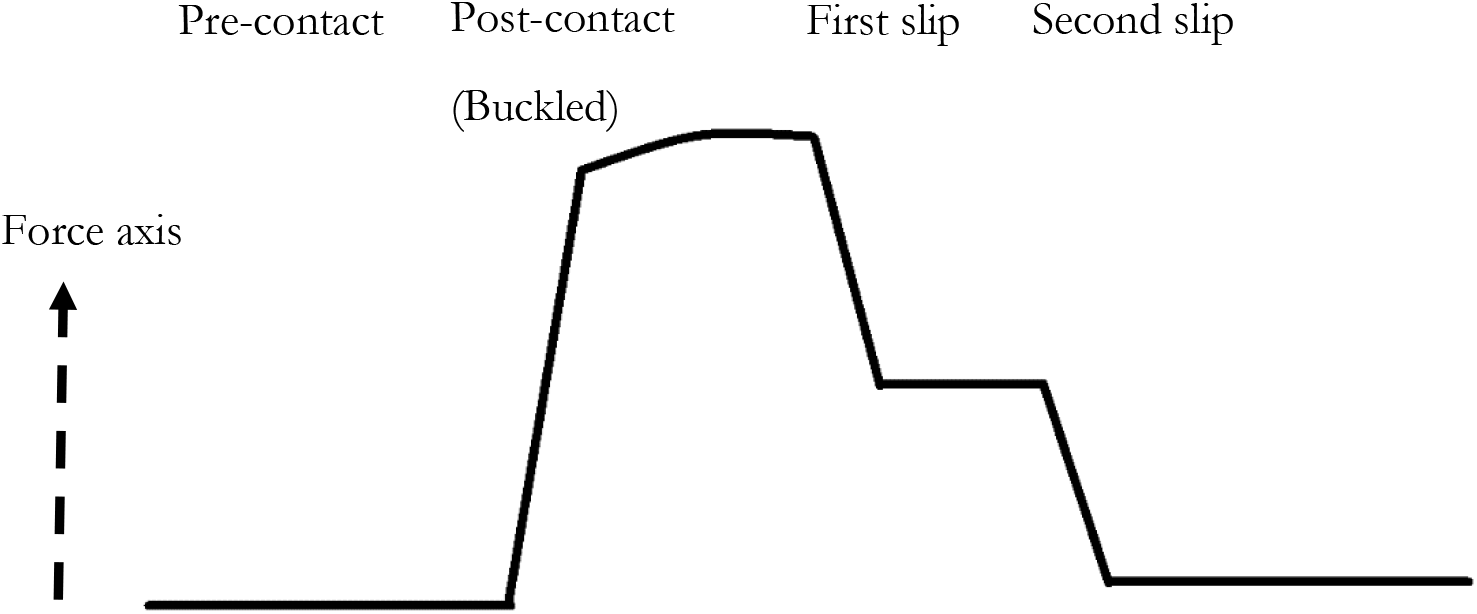
Representation of a typical force vs distance (of headstand) graph of a compliant probe buckling profile with two shanks where slipping of each shank occurs in term after mode one buckling. A residual force remains even after all probes have slipped.

### Insertion force definition

The insertion force is commonly defined as the force required to puncture the surface of the tissue [7], [13]. In this sense it is the same as the maximum indentation that the probe-tissue system can withstand before puncture. Indentation of materials is dictated by contact mechanics with theoretical approaches such as the Hertz model which are used to describe and calculate the force required to indent a material a certain distance.

### Dimpling definition

Dimpling is defined as the maximum indentation distance experienced by the tissue, occurring at puncture. The insertion force is dependent on many variables: speed of insertion, probe tip shape, cross-sectional area, tissue density and wetness of surface to name a few. Greater insertion forces not only increase the risk of buckling for compliant materialled probes, it also imparts more forces into the neural tissue, increasing the trauma of the insertion procedure. Hence, lower insertion forces are desired[13].

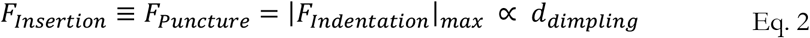

### Shear force definition

The shear force is a contribution of frictional force experienced by the probe on the tissue and the force required to continue the path of the probe in the tissue, the cutting force. Like the insertion force, the shear force is dependent on variables such as insertion speed, cross-sectional area and tissue material. In this paper the shear force is defined as the force experienced by the probe once the probes have been inserted a specified depth, defined here as 1.5 mm since this corresponded to the end of the tapering probes. Speed was not investigated in this study, so a speed that best matches that of surgical insertion is used, 0.01 mm/s.

## Experimental

### Polymer and analysis tools

The most common polymer probes are made from SU-8, polyamide and Parylene C. This investigation uses Parylene C, produced by the Biomedical Microsystems Laboratory at the University of Southern California, but the approaches are equally valid for other polymer probes. Post-data processing was carried out using Origin Software and FEA using COMSOL

### Force testing

An Instron (5940 Series Single Column Tabletop) with a 50g load cell was used to actuate and record all force testing. Forces were recorded as a function of stage displacement. Parylene devices of different shank numbers (1, 2, 4, 6, 8, 16, 24, 32, 48 and 64) and probe lengths (2.75 mm or 5.5 mm) were driven vertically at a constant speed of 0.01mm/s either into a metal plate or the 0.6wt.% agarose phantom for buckling and insertion testing. Before each test, the Instron was calibrated to account for inertial forces. An insertion speed of 0.01 mm/s was chosen to represent a realistic speed at which insertion occurs during surgery and reportedly reduces the stress on the brain[14].

### Agarose

The 0.6 wt. % agarose gel was formulated by dissolving 0.006 g of agarose powder (Sigma-Aldrich) to every 1 mL of deionized water at an elevated temperature. Overnight gelling in a −4 °C fridge ensured full crosslinking occurred to best match the mechanical property of brain tissue[15].

## Results

### Buckling

#### Short vs long probes

As shown in the Table 2, A comparison between the buckling force collected from experimental testing with theoretical values (both the analytical calculations using Euler’s Formulae, which was modified to account for tapering and finite element analysis using COMSOL) was made (Figure 3). A good agreement for the 5.5 mm probes, between the theoretical and the collected data was found. Theoretical values seem to slightly overestimate buckling. For short probes the buckling force for a single probe is around 0.57 mN. Theoretical values seem to slightly overestimate buckling. Most 5.5 mm probes were unable to insert into 0.6 wt.% agarose.

**Table 1:** Geometry of probes. Simple polymer probes were used, not metalized probes.

|  | Parylene C<br>probe |
| --- | --- |
| *Shanks* | 8 |
| *Thickness* | 20 $\mu\text{m}$ |
| *Width* | 110 - 150 $\mu\text{m}$ |
| *Shank pitch* | 250 $\mu\text{m}$ |
| *Length* | 5.5 mm |
| *Tip angle* | $\sim 45^\circ$ |

**Table 2:** Buckling force for a single Parylene shank at two lengths, 5.5 mm (long) (N=3). Standard deviation is reported as the error.

| Method | Buckled force<br>per shank (mN) | Buckle force per<br>shank (mN) |
| --- | --- | --- |
|  | Long probe | Short probe |
| *COMSOL* | 0.17 | 0.57 |
| *Analytical*<br>*(weighted)* | 0.18 | 0.59 |
| *Experimental* | $0.15 \pm 0.01$ | $0.57 \pm 0.17$ |

#### Does buckling scale linearly with shank numbers?

As shown in the Table 3, shanks converged to within 0.55 ± 0.11 mN for different shank numbers, indicating that the buckling force scales linearly, as expected from theory. Standard deviation is reported as the error (N=3).

**Table 3:** The normalized force per shank for short arrays of different shank numbers was experimentally determined to test the scaling ability of more dense arrays. (N=3).

|  | Shank<br>number | Buckled Force per shank<br>(mN) |
| --- | --- | --- |
| **2D** | 1 | $0.57 \pm 0.17$ |
| | 2 | $0.58 \pm 0.17$ |
| | 8 | $0.55 \pm 0.08$ |
| **3D** | 32 | $0.51 \pm 0.14$ |
| | 48 | $0.60 \pm 0.02$ |
| | 64 | $0.47 \pm 0.09$ |

#### Modelling environment

**Figure 3:**
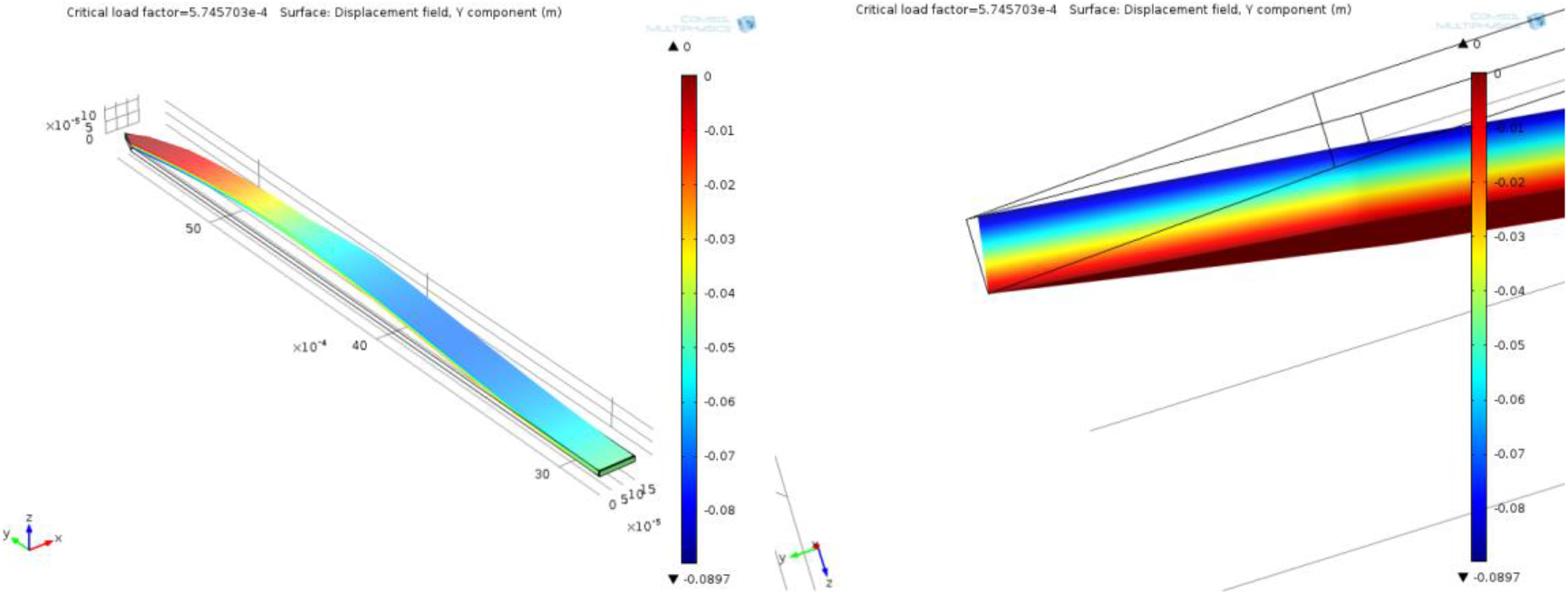
Exact geometry of the long and short long probes were modelled under a linear bulking regime in COMSOL. Mode one buckling for the short probe is depicted. The maximum displacement amplitude is shifted towards the tapered tip end because of the non-uniform width near the tip. A closeup of the tip acting on the ‘metal plate’ demonstrates the non-adhesive contact behaviours and highlights the potential risk of slipping.

### Normalized insertion

#### Do insertion parameters scale linearly with shank number?

The insertion measurements were far harder to measure since the brain phantom introduced an additional layer of error to the results, see table 4.

**Table 4:** Normalized insertion force of short probes into 0.6 wt.% agarose brain phantom. The average insertion force is around 0.18 ± 0.06 mN per shank, suggesting that shank numbers are roughly linearly related to the insertion force. This is around 1/3 of the buckling force for the same probes, satisfying Regime 1 (see theory section). The shear force was around 0.34 ± 0.11 mN per shank, displaying a very rough linear proportionality with shank number too. The 2D data has greater error on their measurements which could be due to their small values. However, the data does seem to be grouped together into 2D and 3D values, probably highlighting different behaviour of the two structures during insertion. (N=3).

|  | Shanks | Normalized<br>insertion force<br>(mN) | Normalized<br>shear force<br>(mN) |
| --- | --- | --- | --- |
| **2D** | 1 | $0.24 \pm 0.09$ | $0.27 \pm 0.11$ |
| | 2 | $0.23 \pm 0.09$ | $0.32 \pm 0.13$ |
| | 4 | $0.23 \pm 0.11$ | $0.33 \pm 0.15$ |
| | 6 | $0.21 \pm 0.09$ | $0.34 \pm 0.16$ |
| | 8 | $0.19 \pm 0.08$ | $0.45 \pm 0.29$ |
| **3D** | 16 | $0.16 \pm 0.02$ | $0.31 \pm 0.04$ |
| | 24 | $0.16 \pm 0.03$ | $0.27 \pm 0.06$ |
| | 32 | $0.15 \pm 0.01$ | $0.26 \pm 0.04$ |
| | 48 | $0.11 \pm 0.01$ | $0.24 \pm 0.03$ |
| | 64 | $0.15 \pm 0.03$ | $0.23 \pm 0.02$ |

## Discussion

### Linear scaling of buckling with shank number?

According to Figure 4, the premise of proportional scaling of forces and dimpling is demonstrated for higher shank numbers. Shear forces were defined as forces measured on the probe at a depth of 1.5 mm. The data follows two separate linear trends, one for 2D arrays (8 shanks or less) and one for 3D.

From table 1, a single short shank, of 2.75 mm length, appears to contribute a single buckling load of around 0.57 mN according to both theoretical (COMSOL & analytical) and experimental data and normalized long probes generate a force of around 0.15 mN. Experimental values indicate that larger shank numbers contributions scale linearly with increasing the number of probes, see table 2, suggesting an average buckling load of 0.55 mN for short probes. This scaling can be explained with the analytical model. Euler’s buckling formula in an altered formant is printed below (Eq. 3).

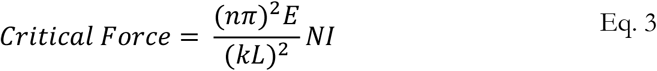

The second moment of area is taken to the side for illustration with an added term *N* representing the number of shanks supporting the force load. All other terms are described in the introduction section. If all terms are held constant while only changing the value for *N*, the results for the critical load would form a straight line passing through the origin, supporting the idea of linear buckling force added up from each additional shank.

### Buckling - short vs long probes

The buckling force differences for a single shank with short and long length should be a multiple of 4 according to the 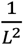 factor (Eq. 3). However, tapering of the probes tip for the short probe will dominate more than the same tapering for the long probes, so the actual second moment of area, I, for shorter probes will be slightly less. The ratio of short to long probes would therefore be expected to be slightly less than 4. From the experimental data, the difference ratio between short and long a single shank is around 3.8. This is expected and indicates that the boundary conditions for both long and short probes are equivalent since boundary conditions are captured by the value k. Asymmetry of the probes profile will again be more dominant in the short probes and likely affected the buckling force via the action of eccentricity.

### Theoretical comparisons

Theoretical values appear to overestimate the buckling force slightly. The experimental probes may suffer from: not being perfectly straight, initial stress in the probes derived from the fabrication process and the presence of friction. These were not accounted for when calculating the theoretical values. The speed of buckling did not affect the buckling force, but faster speeds did increase the risk of the probes slipping before buckling [26]. It is recommended for compression testing of compliant probes to use lower speeds [26]. Slipping of a shank results in other shanks bearing the brunt of the axially placed load, lowering the value for *N* in the modified equation by one, lowering the force required to buckle the probe. This slipping is a feature of the mechanical test setup and not highly representative of buckling on the brain, since dimpling would act as a barrier to slipping for perpendicular insertion at least.

### Are insertion parameters linear with shank number?

The aim of the insertion study was to quantify the forces exerted by the Parylene probes on the brain phantom. From the experimental data, an increase of the number of shanks, meaning an increase in the area or the perimeter of the probes results in a roughly proportional increase of the insertion force, dimpling distance and shear force. In all cases, a linear trend holds rudimentarily for the 2D and 3D probes. However, the trend is slightly steeper for the 2D than for 3D arrays for all parameters measured, suggesting that 3D probes insert differently. Although, this is only slightly notable for the insertion force and dimpling distance, it is quite notable for the shear force.

### Analysis of insertion forces

#### Insertion force

From the experimental data, an increase of the number of shanks, meaning an increase in the area or the perimeter of the probes results in a roughly proportional increase of the insertion force, dimpling distance and shear force Normalizing the insertion force by the number of shanks across 2D and 3D data generates a rough value of 0.18 mN per shank. Comparing this with the normalized short and long shank buckling forces of 0.57 and 0.15 mN respectively, it is clear that regime 1 occurs for the short probes (*F*_*Euler*_ ≫ *F*_*Insertion*_) and regime 2 for the long probes (*F*_*Euler*_ ≤ *F*_*Insertion*_) when inserting into 0.6 wt% agarose brain phantom. However, there seems to be grouping of the insertion data between 2D and 3D devices. For example, the 2D insertion force average was 0.22 ± 0.09 mN whereas 3D was 0.14 ± 0.02 mN.

#### Shear force

The normalized shear force is evaluated to be roughly 0.30 ± 0.11 mN across the 2D and 3D data. Since these were taken at 1.5 mm depth into the gel, only 1.25 mm of probe would be exposed at this point. Correcting for the change of length for the buckled experimental data for both short and long probes, the buckling force should be 4.8 times larger or 2.76 mN and 0.72 mN for short and long probes respectively. Both would satisfy regime 1, although the longer probe is more likely to fall into regime 2, and buckle after partial insertion. However, shear force has two groupings: 2D at 0.34 ± 0.17 mN and 3D at 0.26 ± 0.04 mN like that of the insertion force data.

#### Reasons for 2D and 3D data differences

One reason for the differences is small misalignments when hand stacking the arrays to make the 3D probes. The gel dimples under load from a single probe. If more closely spaced shanks are to be inserted as well, then a slight axial misalignment would change the point at which the nearest neighbour shank would encounter the gel surface. This nearest neighbour shank would either indent the now stressed gel or fall into the dimple, colliding with the leading shank. Both situations would slightly lower the force of insertion and dimpling distance of the probes overall, since the gel is either being held taught or already being cut by the leading probe. 3D probes will be more susceptible to this bending effect since their nearest neighbour will be in the structurally weakest direction, a vector along its thickness.

Shear measurements would suffer from the twin effect of shanks in contact due to initial insertion described above, but also by the probes bending in the gel as it passes through it. The bending would occur in the direction of their thinnest and structurally weakest direction. Such probes would collide with its nearest, bending neighbour. This suggests the effect of changing surface area rather than cross-sectional area plays a more important role on shear force. The friction contribution would be lower for shanks with one of their faces in contact. The cutting contribution would also be lower since it is likely that the probes in contact will be staggered, with one probe creating the path through the gel and the second running along the shank-gel interface. This probable explains the shallow gradient for the 3D probes for each parameter, especially the shear measurements.

The theoretical regimes and the behaviour observed from the data match, confirming a link between the measured forces of buckling, insertion and shear. There is much greater variability of insertion and shear forces measured for different shank numbers than for their buckled forces, making this insertion normalization a rule of thumb approach only for increased shank numbers.

#### Improvements for future insertion testing

It is suggested that for future 3D stacking of compliant shanks, that the separation distances along the stacking direction should be larger than their 2D separation distance. Rigid 3D probes one to one ratio of 2D:3D separation distance does not appear to hold for compliant materials. This paper only looked at 3D grid configuration of shank stacking. The effect of staggering one layer from the next, whilst keeping the same separation distance might alleviate the issues of shank collision. This would also benefit from more evenly covering the same volume of tissue with electrodes. Alternatively, or in addition to, structures that incorporate an arch or ridge at the base of the shanks may help to stiffen the probes in its thickness direction, without increasing the cross-sectional area and hence the trauma.

The probes bending in tissue is related to its second moment of area, I, but trauma is related to the cross-sectional area, A. These two quantities are slightly different for the rectangular shanks discussed in this paper, see the formulas below.

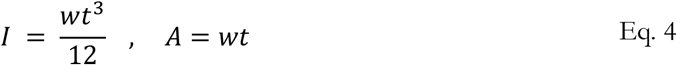

Where w is the width and t the thickness. For the same cross-sectional area as the probes used in this study, a greater second moment of area could be produced by making a square cross-section. This would also reduce the surface area presented to the gel or tissue as the shanks are being inserted through the material, likely lowering the shear force exerted on the tissue and possibly reducing bending forces acting on the shank itself. It has been reported that asymmetry in needles results in bending. The one-sided tapering of the Parylene probes might be exacerbating the bending forces experienced in the gel.

One workaround for the bending of shanks on initial insertion would be to extend the PEG coating for more than half the length of the probes, exposing only the tips. It should be noted that the brain does not present a uniform surface to oncoming probes. During surgical testing, such probes will encounter the tissue at slightly different times, even if the probes are perfectly aligned. Increasing the stiffness in the weakest direction by the methods above, increasing the 3D spacing and encasing more of the shanks length in PEG should ensure that less shanks should collide during insertion, and then the gradient of the 2D and 3D should be equivalent.

#### Testing errors

Minor misalignment of the probes during buckling testing would also affect the results. There are two main sources of misalignment. The first is loading of the probes on to the Instron. If the probe that were mounted on the clamps were partially tilted, the shanks would encounter the metal surface at slightly different times. The second is a potential consequence of the handcrafted 3D stacking method. If different layers of arrays were stacked at slightly different lengths, different contact times may be experienced by the shanks and different associated buckling forces for those probes. These are unlikely to play a major role since probes were examined carefully when stacking and placing on the Instron. These minor misalignments may explain the slight differences in normalized forces for larger stacked arrays.

## Conclusion

Compliant materials show promise in improving the long-term stability of neural probes inside the body but are limited by their tendency to buckle. It was shown that a buckling force of around 3x the insertion force results in consistent successful insertion, a useful rule of thumb when designing neural probes for applications. The forces of import for neural probes during insertion were shown, using normalized analysis, to be roughly linearly related to the number of shanks in a device, although different behaviour of 2D and 3D probes were observed. The issue of slipping was addressed when mechanically buckle testing compliant or flexible probes by reducing the headstand speed. One long-term goal for neural probes is to make them thinner, reducing trauma and improving the resolution of information from the brain. However, this presents a problem as this greatly reduces the buckling strength of the probes. More dense arrays with greater shank number will increase the buckling force. Since most techniques test on single shanks, not dense arrays and most goals specify dense arrays, the problem could be less of an issue than previously thought.

## Acknowledgements

I would like to thank the Charles Vest International Scholarship for sponsoring my research visit at the University of Southern California and the Biomedical Microsystems Laboratory for hosting me.

## References

[1] J. Subbaroyan, D. C. Martin, and D. R. Kipke, ‘A finite-element model of the mechanical effects of implantable microelectrodes in the cerebral cortex’, J. Neural Eng., vol. 2, no. 4, pp. 103–113, Oct. 2005.

[2] V. S. Polikov, P. A. Tresco, and W. M. Reichert, ‘Response of brain tissue to chronically implanted neural electrodes’, J. Neurosci. Methods, vol. 148, no. 1, pp. 1–18, Oct. 2005.

[3] T. D. Y. Kozai, A. S. Jaquins-Gerstl, A. L. Vazquez, A. C. Michael, and X. T. Cui, ‘Brain Tissue Responses to Neural Implants Impact Signal Sensitivity and Intervention Strategies’, ACS Chem. Neurosci., vol. 6, no. 1, pp. 48–67, Jan. 2015.

[4] R. Biran, D. C. Martin, and P. A. Tresco, ‘Neuronal cell loss accompanies the brain tissue response to chronically implanted silicon microelectrode arrays’, Exp. Neurol., vol. 195, no. 1, pp. 115–126, Sep. 2005.

[5] T. D. Y. Kozai, A. L. Vazquez, C. L. Weaver, S.-G. Kim, and X. T. Cui, ‘In vivotwo-photon microscopy reveals immediate microglial reaction to implantation of microelectrode through extension of processes’, J. Neural Eng., vol. 9, no. 6, p. 066001, Oct. 2012.

[6] J. C. Williams, R. L. Rennaker, and D. R. Kipke, ‘Long-term neural recording characteristics of wire microelectrode arrays implanted in cerebral cortex’, Brain Res. Protoc., vol. 4, no. 3, pp. 303–313, Dec. 1999.

[7] W. Jensen, U. G. Hofmann, and K. Yoshida, ‘Assessment of subdural insertion force of single-tine microelectrodes in rat cerebral cortex’, in Proceedings of the 25th Annual International Conference of the IEEE Engineering in Medicine and Biology Society (IEEE Cat. No.03CH37439), 2003, vol. 3, pp. 2168–2171 Vol.3.

[8] M. Jeon et al., ‘Partially compliant MEMS neural probe composed of polyimide and sucrose gel for reducing brain damage during and after implantation’, J. Micromechanics Microengineering, vol. 24, no. 2, p. 025010, Jan. 2014.

[9] J. Agorelius, F. Tsanakalis, A. Friberg, P. T. Thorbergsson, L. M. E. Pettersson, and J. Schouenborg, ‘An array of highly compliant electrodes with a tailored configuration locked by gelatin during implantation—initial evaluation in cortex cerebri of awake rats’, Front. Neurosci., vol. 9, 2015.

[10] M. A. Howard, B. A. Abkes, M. C. Ollendieck, M. D. Noh, C. Ritter, and G. T. Gillies, ‘Measurement of the force required to move a neurosurgical probe through in vivo human brain tissue’, IEEE Trans. Biomed. Eng., vol. 46, no. 7, pp. 891–894, Jul. 1999.

[11] E. Fernández et al., ‘Acute human brain responses to intracortical microelectrode arrays: challenges and future prospects’, Front. Neuroengineering, vol. 7, 2014.

[12] G. Kook, S. W. Lee, H. C. Lee, I.-J. Cho, and H. J. Lee, ‘Neural Probes for Chronic Applications’, Micromachines, vol. 7, no. 10, p. 179, Oct. 2016.

[13] P. J. Gilgunn et al., ‘An ultra-compliant, scalable neural probe with molded biodissolvable delivery vehicle’, in 2012 IEEE 25th International Conference on Micro Electro Mechanical Systems (MEMS), 2012, pp. 56–59.

[14] D. Lewitus, K. L. Smith, W. Shain, and J. Kohn, ‘Ultrafast resorbing polymers for use as carriers for cortical neural probes’, Acta Biomater., vol. 7, no. 6, pp. 2483–2491, Jun. 2011.

[15] S. Takeuchi, D. Ziegler, Y. Yoshida, K. Mabuchi, and T. Suzuki, ‘Parylene compliant neural probes integrated with microfluidic channels’, Lab. Chip, vol. 5, no. 5, pp. 519–523, Apr. 2005.

